# A voltage-based metaplasticity rule applied to the model hippocampal granule cell accounts for homeostatic heterosynaptic plasticity

**DOI:** 10.1101/557173

**Authors:** Azam Shirrafiardekani, Lubica Benuskova, Jörg Frauendiener

## Abstract

Long-term potentiation (LTP) and long-term depression (LTD) of synaptic efficacies are involved in establishment of long-term memories. In this process, neurons need to adjust the overall efficacy of their synapses by using mechanisms of homeostatic plasticity to balance their activity and control their firing rate. For instance, in the dentate granule cell *in vivo,* induction of homosynaptic LTP in the tetanized medial perforant path is accompanied by heterosynaptic LTD in the non-tetanized lateral perforant path. We used the compartmental model of this cell to test the following hypotheses: 1. Using plasticity and metaplasticity rules both based on postsynaptic voltage we can reproduce homosynaptic LTP and concurrent heterosynaptic LTD, provided there is an ongoing noisy spontaneous activity; 2. Frequency of an ongoing noisy spontaneous activity along the lateral path determines the magnitude of heterosynaptic LTD. In experiments where procaine was used to block the lateral spontaneous activity, no heterosynaptic LTD occurred. However, when the procaine was washed out and a second tetanization was applied to the medial path, no heterosynaptic LTD could have been induced neither. Our simulations predict that the reduced frequency of spontaneous activity in the lateral perforant path can account for this lasting absence of heterosynaptic LTD.

## 1. Introduction

Although during the last century our understanding of how neurons work and how they process information has grown rapidly, we have still not managed to find a comprehensive theory to explain memory and learning phenomena. In fact, experimental studies of a variety of brain regions reveal that several mechanisms may be involved in memory formation and learning. The common denominator is the ability of synapses to change their efficacy in an activity-dependent way, which is referred to as synaptic plasticity. Neural stem cells are able to generate new neurons in the process called neurogenesis and new investigations show that neurogenesis of the dentate gyrus might be involved in the mechanism of memory encoding (Deng et al., 2010). However, it is not clear how these newborn neurons engage in existing memory circuits even when synaptic plasticity is involved. An investigation by Snyder et al. (2001) supports the idea that synaptic plasticity of adult-born dentate granule cells is involved greatly in the process of learning and memory formation. Because of the crucial role of synaptic plasticity in learning and memory, in this paper we investigate phenomena of synaptic plasticity in the adult dentate granule cell. Two forms of long-lasting synaptic plasticity are long-term potentiation (LTP) and long-term depression (LTD). These can occur simultaneously in neighboring synaptic pathways onto dentate granule cells. LTP and LTD can be induced homosynaptically or heterosynaptically. Homosynaptic plasticity occurs when synapses are tetanized directly by high-frequency presynaptic stimulation (HFS) and heterosynaptic plasticity occurs on the untetanized synapses. According to several experimental studies, heterosynaptic plasticity mechanisms are necessary for complementing the homosynaptic plasticity during the homeostatic regulation of neuronal activities (Chen et al., 2013). In the dentate granule cells, when the medial perforant path (MPP) is tetanized by high frequency stimulation (HFS), homosynaptic LTP occurs in this path and concurrently heterosynaptic LTD occurs on the neighboring lateral perforant path (LPP) (Douglas and Goddard, 1975; Levy and Steward, 1979; Abraham and Goddard, 1983; Doyère et al., 1997). Because of this interesting property of granule cells, our investigation of synaptic plasticity is based on this particular cell type. In this work, we used computer simulation to test several hypotheses related to synaptic plasticity induction. One of our assumed mechanisms is based on the nearest-neighbor pair-based STDP (spike timing-dependent plasticity), as experimental studies have shown that the precise timing between pre- and postsynaptic spikes can play a role in LTP and LTD induction (Markram et al., 1997). Investigations by Lin et al. (2006) support a role for STDP in granule cell plasticity, using pre- and postsynaptic spikes paired in either pre-post or post-pre order. Furthermore, a plasticity model is more plausible if each presynaptic spike pairs only with the two most recent postsynaptic spikes; this mechanism is known as nearest-neighbor STDP (Izhikevich and Desai, 2003). However, in this study we replace the postsynaptic spike with postsynaptic voltage as other studies bring evidence that it is the postsynaptic voltage and not the postsynaptic spikes that is crucial for induction of synaptic plasticity. Thus, the STDP becomes rather an ETDP, i.e. event timing-dependent plasticity. Another mechanism that is lately assumed to be involved in synaptic plasticity induction is metaplasticity, defined as the ability of previous activity of a neuron to regulate subsequent synaptic plasticity. Metaplasticity can act homeostatically to protect synaptic weights from extreme increases or decreases (El Boustani et al., 2012). Having such a homeostatic mechanism is necessary to regulate the neural firing rate and adjust to any other neural activities that may destabilize the neural networks. The third factor we include in our model is an ongoing spontaneous activity of neural circuits *in vivo.* For instance, the prominent state of *in vivo* hippocampal network is theta activity (Buzsaki 2002). However, in theory and modelling studies, only in the Bienenstock, Cooper and Munro (BCM) theory of synaptic plasticity (Bienenstock et al., 1982; Cooper et al., 2004) the role of spontaneous activity in the outcome of monocular deprivation was taken into account for the first time. In our previous work by (Benuskova and Abraham 2007; Jedlicka et al. 2015, Shirrafiardekani, 2015), the role of spontaneous neural activity in controlling the dynamics of homo- and heterosynaptic plasticity in the dentate gyrus was corroborated. A prediction was made that if the spontaneous activity on the LPP would be blocked then no heterosynaptic LTD would occur. This prediction follows from the point model of a neuron without dendrites where STDP and metaplasticity based on the postsynaptic spike count were used (Benuskova and Abraham, 2007). The same prediction also follows from the multi-compartmental model of granule cell (Jedlicka et al. 2015, Shirrafiardekani, 2015), endowed with the “event timing-dependent synaptic plasticity” based on postsynaptic voltage and metaplasticity based on the postsynaptic spike count (Jedlicka et al. 2015) or the somatic postsynaptic voltage (Shirrafiardekani, 2015).

In our previous work (Benuskova and Abraham 2007; Jedlicka et al., 2015), the metaplasticity rule was based on a postsynaptic spike count. In this paper, however, we use a modified metaplasticity rule, which is based on the postsynaptic voltage rather than the postsynaptic spike count, which we think might be more realistic account, because it does not depend on whether the postsynaptic cell generates spikes or not. We explore two hypotheses by combining ETDP and metaplasticity rules accompanied by noisy spontaneous activity in the compartmental model of a granule cell (GC). The hypotheses we addressed were:

1. The synaptic plasticity and metaplasticity rules based solely on postsynaptic voltage can replicate homosynaptic LTP in the tetanized path and concurrent heterosynaptic LTD in the neighboring non-tetanized path for a variety of protocols.
2. The frequency of noisy spontaneous activity along the LPP determines the magnitude of heterosynaptic LTD when the medial LTP occurs.

To test these hypotheses and examine our (meta)plasticity model we simulated one experimental protocol from Abraham et al. (2001) and several different experimental protocols from Abraham et al. (2007). The first experimental protocol consisted of simulation of application of HFS to MPP to investigate induction of homosynaptic LTP in this path and concurrently heterosynaptic LTD in the neighboring lateral perforant path (LPP).

In the second simulation, we replicated data from Abraham et al. (2007), in which HFS was again applied to the medial path. However, in this protocol, the experimenters blocked the spontaneous activity in the LPP by procaine during the duration of medial HFS. To simulate the inhibition effect of the procaine on the lateral path spontaneous activity, the simulated presynaptic spontaneous activity was switched off during the medial HFS. As in the real experiment, also in our simulations, no heterosynaptic LTD in the LPP accompanied medial LTP.

In the third protocol, two HFS following each other were applied to the medial path. After applying the first HFS, the second HFS with the same pattern as the first one was applied to the medial path. In this simulation, we examined the plasticity impact of the first medial HFS on the synaptic plasticity caused by the second medial HFS. Both medial HFS led to homosynaptic LTP in this path and concurrently heterosynaptic LTD in the neighboring lateral perforant path (LPP).

The aim of the fourth simulation was to examine the effect of procaine inhibition of the lateral spontaneous activity during the first HFS upon the outcome of the second HFS. Thus, two HFS following each other were applied to the medial path except that the spontaneous activity was switched off in the LPP during the first medial HFS only. Then it was turned on again. As before, no heterosynaptic LTD in the neighboring LPP occurred during the first HFS when LPP spontaneous activity was off. Surprisingly, no heterosynaptic LTD occurred during the second HFS even though the lateral spontaneous activity has resumed.

## 2. Methods

### 2.1 Modeling the dentate granule cell

In our simulations, we used the NEURON simulation environment, version 7.3 (Carnevale and Hines, 2006). For the reduced-morphology multi-compartmental model of GC, each section represents a portion of dendritic tree with different lengths, diameters and ion channel densities. The reduced-morphology model has two dendritic branches and a soma, each dendrite contains four sections and the soma contains one section while each section contains seven ion channels described by (Aradi and Holmes, 1999) with parameters as listed in Santhakumar et al (2005). The GC files can be downloaded from the ModelDB at http://senselab.med.yale.edu/modeldb/, accession No. 51781. As we were interested only in the generation of action potentials and not in their propagation, we did not include the axon in our GC model. We used nine compartments in order to reduce the simulation time. We also did our simulations with 29 compartments (14 compartments for middle and distal dendrites in the two main branches and one compartment for the soma) and did not see any significant change in our results. Seven ion channels are implemented in this model: fast sodium (Na), fast delayed rectifier potassium (fKDR), slow delayed rectifier potassium (sKDR), A-type potassium (KA), large conductance and T-type (TCa), N-type (NCa), and L-type (LCa) calcium channels. We also did our simulations with nine ion channels which included calcium-voltage-dependent potassium (BK) and small conductance calcium-dependent potassium (SK) channels, but the results were very close to when only seven ion channels were implemented. Figure 2.1 shows a schematic picture of the cylindrical 9-compartmental model of the dentate granule cell. The granule cell layer (GCL) is the closest dendritic part to the soma. Proximal dendrites (PD), middle dendrites (MD) and distal dendrites (DD) are the second, third and farthest dendritic compartments to the soma, respectively. The granule cell receives two main excitatory inputs (there are no inhibitory synapses in our model) on the middle and distal compartments of the dendrites and learning occurs on all excitatory synapses. Presynaptic spikes at MPP and LPP axons cause postsynaptic potentials in the GC dendrites, which propagate in all directions, while action potentials generated in the soma only propagate back into the dendrites since we do not simulate the axon. The MPP and LPP have 150 synapses each; each dendrite has 75 MPP synapses and 75 LPP synapses evenly distributed along the MD and DD, respectively.

**Figure 2.1:**
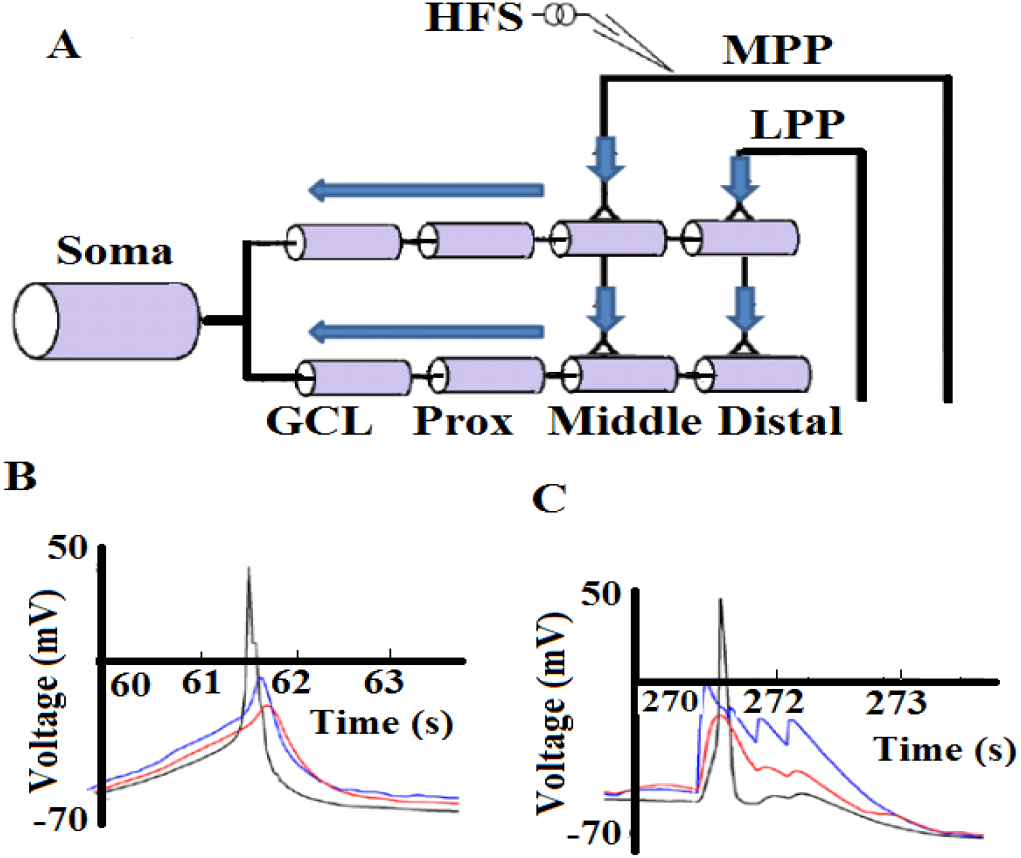
(A) A cylindrical 9-compartmental model of the dentate GC has two dendrites with 300 excitatory synapses (each dendrite has 150 synapses). Each dendrite consists of four compartments and soma has one compartment. This model does not have an axon. The lateral perforant pathway (LPP) relays presynaptic inputs from the entorhinal cortex to the distal dendrites (DD), and medial perforant pathway (MPP) relays presynaptic inputs from the entorhinal cortex to the middle dendrites (MD). Black input lines showing LPP and MPP pathways are only for illustration, because input spikes are delivered directly to synapses in our model. Filled arrows show the flow of input activities towards soma where they may or may not result in generation of an action potential. (B) Somatic action potential (black) and backpropagation of action potential plus EPSPs from MPP (blue) and LPP (red) before HFS. (C) Somatic membrane potential (black) and backpropagation of action potential plus EPSPs from MPP (blue) and LPP (red) during one train of HFS delivered to MPP.

In our simulations, equations were solved with the Crank-Nicholson method. We use this method because it is 2^nd^ order accurate for small time steps. The error in Crank-Nicholson method is proportional to d*t*^2^ and it uses a staggered time step algorithm to avoid iteration of nonlinear equations (Hines and Carnevale, 1997). The time step that we have chosen for the numerical integration is 0.2 ms; we also tested our model with different values of time steps dt = 0.1 ms, 0.07 ms, 0.05 ms, 0.027 ms, 0.026 ms and 0.02 ms and observed the same results.

### 2.2 Simulation of spontaneous presynaptic activity

Our model granule cell receives ongoing presynaptic noisy spontaneous activities via both the medial and lateral pathways from the entorhinal cortex. Spontaneous activity was generated independently and randomly as Poisson spike trains along the MPP and LPP. The Poisson process is a simple yet accurate model of a neuron’s spontaneous firing (Fellous et al., 2003). In the real brain, spontaneous activity is the result of interactions within neural networks and the electrophysiological properties of a single neuron. This activity is related to the functional state of the brain with the key elements of the level of activation of neuromodulatory systems (Herz et al., 2006). In this work, the frequency of presynaptic spontaneous activity has been chosen to be <10 Hz (Gloveli et al., 1997). In our compartmental model of the neuron, presynaptic spontaneous activity is simulated by using independent spike generators (NEURON’s built-in point process NetStim). In the NEURON code, the interspike interval (ISI) of spiking activity is generated according to the following equation:

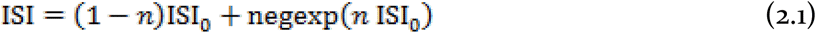

where, *n* is the noise function with 0 < *n* < 1, negexp(*x*) is the negative exponential distribution, which is equal to the homogeneous Poisson distribution with probability of the next spike occurring after time ISI. When *n* is zero, the ISI is equal to ISI_0_ (initial value of ISI) and spiking activity is periodic. When *n* is between zero and one, the spiking activity is quasi-periodic. When *n* = 1, then the spike series obeys the homogeneous Poisson distribution. For all simulations, we have chosen *n* =0.02.

### 2.3 Applying HFS for LTP induction

Another source of input activity that was represented in experiments was tetanization by high frequency stimulation (HFS). In all simulations, HFS was applied to the medial perforant path (MPP) along with an ongoing noisy spontaneous presynaptic activity as described above. A pattern of HFS was the so-called 400 Hz DBS (delta burst stimulation). The 400 Hz DBS consists of 5 trains at 1 Hz and each train contains 10 spikes at 400 Hz. Bursts of these five trains are repeated 10 times at every 30 or 60 s (Figure 2.2).

**Figure 2.2:**
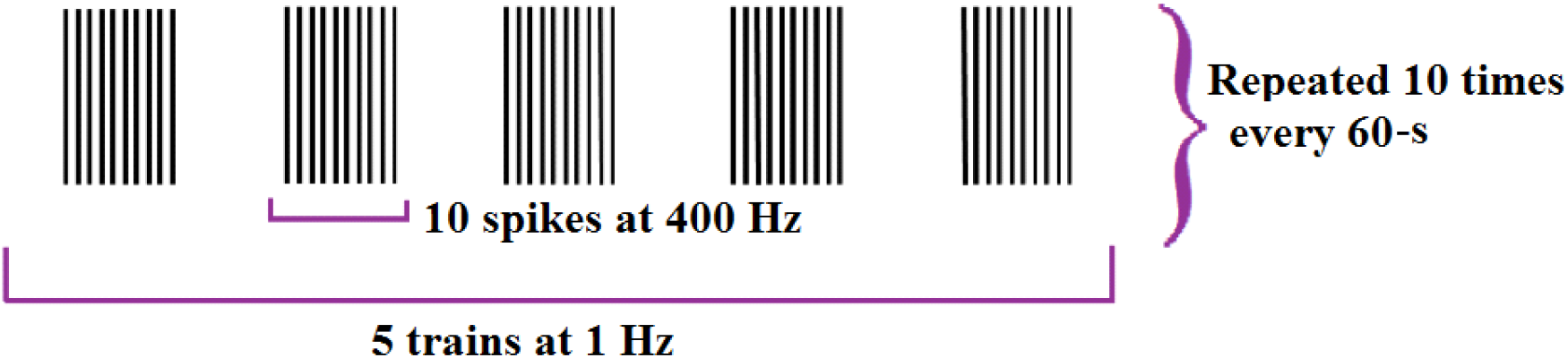
Schematic illustration of the 400 Hz delta burst stimulation (DBS) form of HFS, which consists of 5 trains at 1 Hz, while each train contains 10 spikes at 400 Hz. Bursts of these five trains are repeated 10 times at every 30 or 60 s. HFS was delivered to MPP along with an ongoing noisy spontaneous presynaptic activity as described in the text.

### 2.4 Synaptic plasticity rule

The idea of spike-pair interactions in the dentate granule cell is based on the experimental study of Lin et al. (2006), which supports a role for STDP in granule cell plasticity, based on results of pre- and postsynaptic spikes paired in either pre-post or post-pre order. To model the synaptic plasticity of the dentate granule cell we start with equations for the standard spike-pair STDP (Izhikevich and Desai, 2003)

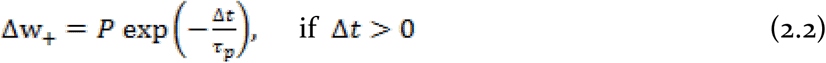

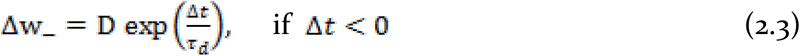

Here Δt= *t_post_* – *t_prs_* is the difference between the arrival time of the post- and presynaptic spikes, respectively. *τ_p_* > o is the decay constant of windows for LTP and *τ_d_* > o is the decay constant of windows for LTD. P > 0 is the amplitude of potentiation, D > 0 is the amplitude of depression. Based on results of Izhikevich and Desai about the relationship between STDP and BCM theory, in order to calculate the overall synaptic change, we used the nearest-neighbor interaction expressed by the following equation:

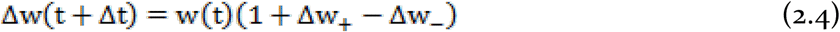

Another reason to use the nearest neighbors is because the back-propagation of postsynaptic spikes to the medial and lateral synapses resets the membrane voltage in the dendritic spines. Therefore, the most recent postsynaptic spikes suppress the effect of previous postsynaptic spikes, which results in the membrane voltage being affected mostly by the recent postsynaptic spike. However, the first difference in implementation in our compartmental model is that *t_post_* is the time when the postsynaptic voltage at the site of a particular synapse crosses certain dendritic voltage threshold. This threshold is the same for synaptic depression and potentiation, and may correspond to a membrane voltage at which the magnesium block is removed from NMDARs, as we know that both LTD and LTP in granule cells are NMDAR-dependent. Based on investigating different values we have arrived at the dendritic threshold value of −37 mV (Jedlicka et al., 2015). In such a way, postsynaptic spikes are replaced by postsynaptic membrane events (Jedlicka et al., 2015). Event timing dependent plasticity happens individually at each synapse as postsynaptic voltage can cross the threshold for an event at different times at each synapses. To calculate the event-timing-dependent plasticity (ETDP) and metaplasticity, additional mod files are implemented into the NEURON code and included in the GC model. Thus, the complete set of simulation files is available for download from the ModelDB database (accession number 185350): http://senselab.med.yale.edu/modeldb/.

### 2.5 Metaplasticity rule

In addition to replacing timing of postsynaptic spikes with timing of postsynaptic voltage events (Jedlicka et al., 2015), Benuskova and Abraham (2007) and Jedlicka et al. (2015) proposed that the amplitude of potentiation P and amplitude of depression D are not constant but rather dynamically change their values as a function of the average of the postsynaptic activity over some recent past. Thus at each time instant an updated value of potentiation and depression amplitudes are calculated as

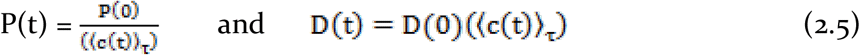

where P(0)>0 and D(0)>0 are positive constants representing initial values of potentiation and depression amplitudes. The temporal average of the postsynaptic activity 〈*c*〉_τ_ is calculated as a running average of the spike count such that:

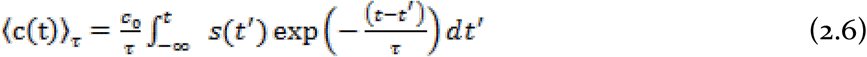

Where s(t’) = 1 if there is a postsynaptic spike at *t*’ or zero otherwise, *C_o_* is the scaling constant and *τ* is the integration period.

In this paper, we introduce a modification, in which the running average of postsynaptic activity is calculated based on the difference between the postsynaptic voltage and resting potential at the soma. Therefore, according to our modification, the average of postsynaptic activity is calculated by the following integral:

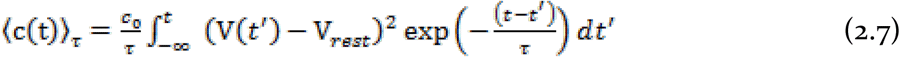

The scaling constant *c*_0_ is equal to 0.0025 mV^-2^. (y(t′) – V_rest_) is the difference between the postsynaptic voltage and resting potential in the soma. V_rest_ is the initial resting potential and equal to −75 mV. By taking the square of the voltage difference, we ensure that (〈c(t)〈_τ_) ≥ 0.

Another synaptic plasticity rule, albeit different from ours, that is based on the postsynaptic voltage is the model introduced by Clopath et al. (2010). In the metaplasticity part of their rule only the depression amplitude depends on the average of postsynaptic voltage while the potentiation amplitude remains constant. To test their idea with our synaptic plasticity model we tried to find out, whether we can observe the same synaptic plasticity results when only one of the potentiation or depression factors is subject to the metaplastic change. Therefore, for our metaplasticity, we ran the simulations with setting the P factor at a constant value and the D factor as a function of the average postsynaptic activity (〈c(t)〈_τ_) i.e.

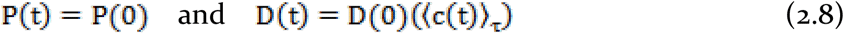

And vice versa

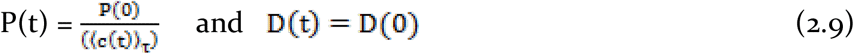

where, P(0)>o and D(0)>o are initial amplitude values for synaptic potentiation and depression, respectively, and (〈c(t)〈_τ_) is the average of postsynaptic activity expressed by equation (2.7).

We tried all scenarios for metaplastic control of potentiation and/or depression amplitudes expressed by equations (2.5), (2.8) and (2.9). In neither case there was a need for a fixed upper and/or lower bound on the weights. We present results for P being metaplastically controlled while the depression factor D being constant, i.e. Equation (2.9).

## 3. Results

### 3.1 First simulation – heterosynaptic plasticity due to a single HFS

We simulated HFS (section 2.3) applied to the medial pathway along with the simulated ongoing spontaneous activity (section 2.2). After applying the medial HFS, homosynaptic LTP and concurrent heterosynaptic LTD occurred in the multi-compartmental model of the dentate granule cell (Figure 3.1) like in (Abraham et al., 2001). Parameter values are taken from Table 3.1.

**Table 3.1.**
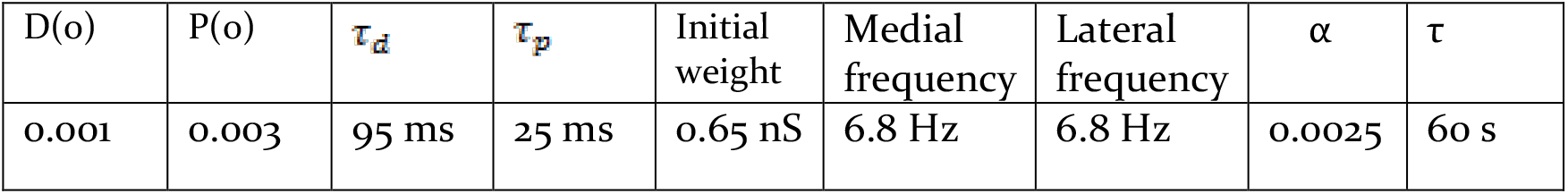
Parameter values for optimal match with the experimental data.

**Figure 3.1:**
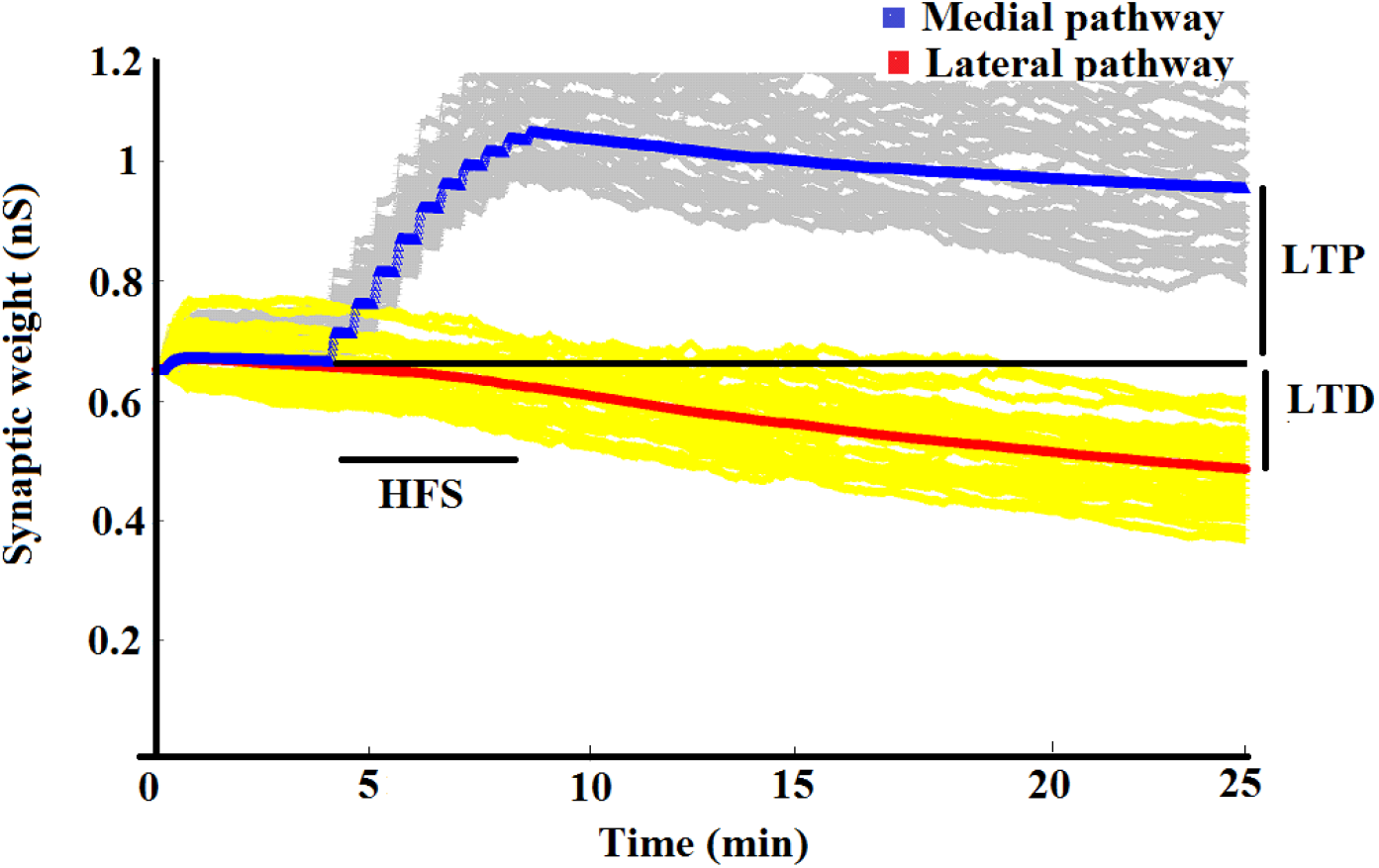
Evolution of synaptic weight expressed in terms of postsynaptic conductance over time. The blue trace corresponds to the average weight of all 150 medial synapses, while the red trace represents the average weight of all 150 lateral synapses. Grey traces represent individual medial synapses and yellow traces represent individual lateral synapses. The simulation is started with the same initial weight value for all synapses that is adjusted so that the granule cell model fires postsynaptic action potentials at around 1 Hz frequency. Before medial HFS, both average weight traces are stable on average, until starting the medial HFS (black horizontal line). During HFS, the synaptic weights increase in the medial pathway and decrease in the lateral pathway. The percentage of LTP and LTD is calculated as the difference between the average weight at 25 minutes from the baseline of the blue curve (for the medial pathway) and from the red curve (for the lateral pathway). In this run, we observe +37% LTP in the medial pathway and −27% LTD in the lateral pathway.

When the simulation starts, after a short while all synaptic weights and the average synaptic weights become stable and stay at the 100% baseline. This stable state lasts until the first burst of medial HFS occurs. When medial HFS starts the average of the medial synaptic weights increases and the average of the lateral synaptic weight decreases from the baseline. To calculate the size of LTP and LTD we measured the magnitude of the medial and lateral synaptic weight as a difference from the 100% baseline level one minute before the first burst of HFS and one minute before the end of the simulation. Therefore, numbers over 100% baseline show the percentage of LTP magnitude and less than 100% show the percentage of LTD magnitude. Our model with parameter values from table 3.1 produced LTP in the medial pathway at approximately 37% from the baseline and LTD in the lateral pathway at approximately −27% from the baseline. In comparison, in the experimental studies (37 ± 5%) LTP in the medial pathway and (30 ± 5%) LTD in the lateral pathway was observed (Abraham et al., 2001). Magnitudes of LTP and LTD as a function of different values of parameters are illustrated in Figure 3.2. In general, all parameter values should be adjusted so that the output frequency of the model GC in response to the spontaneous input activity should be around 1 Hz (Kimura and Pavlides, 2000).

**Figure 3.2:**
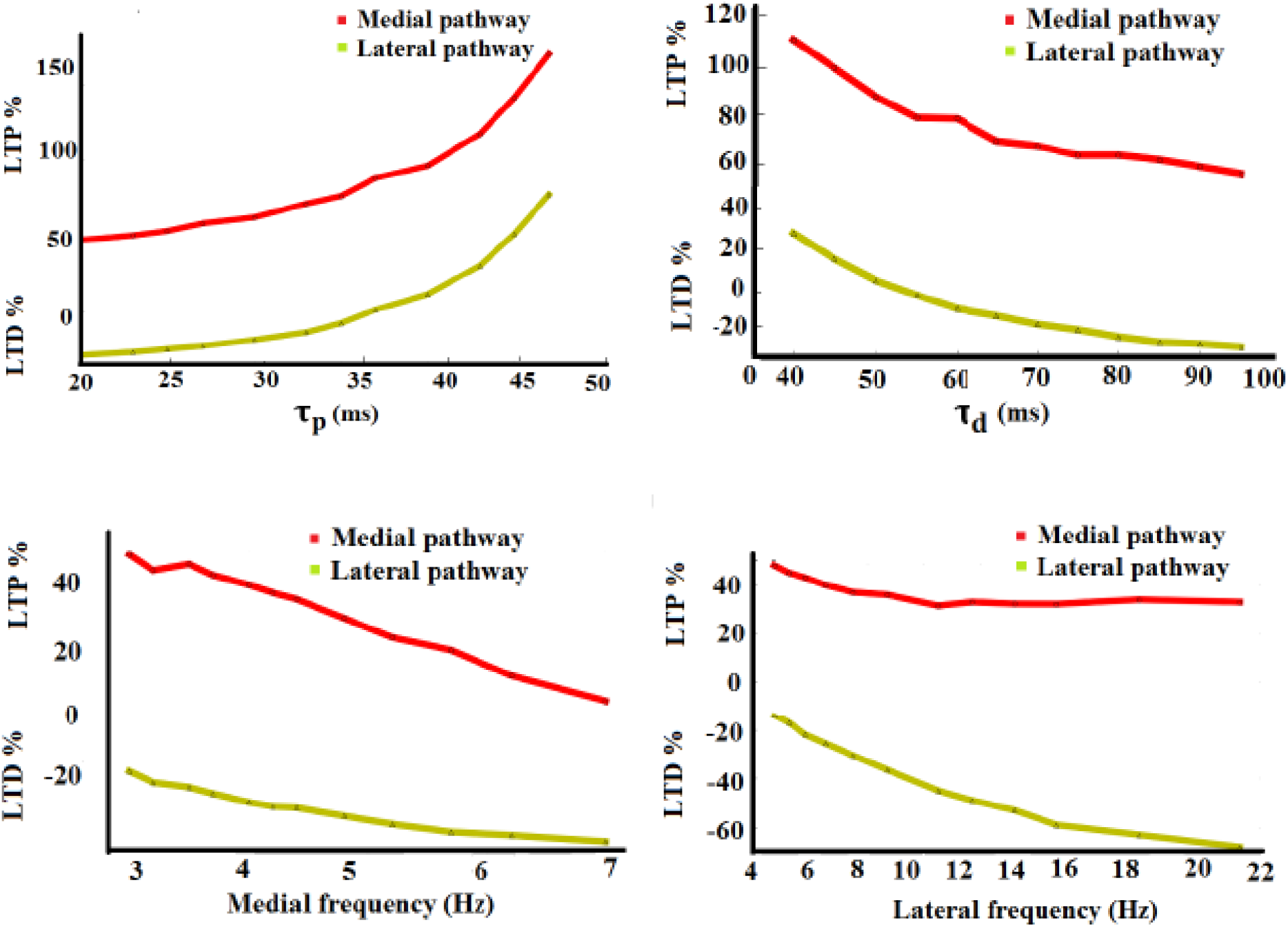
Magnitudes of LTP and LTD for different values of (A) τ_p_, (B) τ_d_, (C) medial spontaneous frequency, and (D) lateral spontaneous frequency.

Thus, we tested and corroborated the hypothesis that the voltage-based plasticity and metaplasticity model with the reduced morphology multi-compartmental model of the GC combined with the noisy spontaneous activity yields homosynaptic LTP and concurrent heterosynaptic LTD.

#### 3.3.1 Detecting pre- and postsynaptic events

In our synaptic plasticity model, we need to detect pre- and postsynaptic events to calculate the nearest-neighbor pre- and postsynaptic event interactions. In our compartmental model of granule cell, the presynaptic spikes are presynaptic events, which are generated at the location of each synapse and lead to the generation of the excitatory postsynaptic potential (EPSPs) at the corresponding site of the dendrite. From there, EPSPs propagate along the dendrites in all directions because of the action of ion channels in the membrane and because of the passive electric properties of the dendritic model. Now we explain how to compute the postsynaptic event. Once the simulation starts and input activities, either spontaneous or HFS, are delivered to the granule cell, the somatic membrane potential depolarizes, which increases the voltage above the initial resting potential. If the summation of input activities (in the form of EPSPs) is strong enough to depolarize the somatic membrane potential past the firing threshold, the granule cell produces a postsynaptic action potential which propagates back to the dendrites (Mehta, 2004). The instant when the sum of synaptic EPSP and/or bAP exceeds a certain threshold is referred to as the time of a postsynaptic event (Jedlicka et al., 2015). The dendritic threshold for detection of a postsynaptic event is −37 mV (Krueppel et al., 2011). This voltage is critical for induction of synaptic plasticity (Lisman and Spruston, 2005).

In our model, we have only one threshold, which applies to both LTP and LTD. Figure 3.3A shows the membrane voltage with one postsynaptic spike occurring in the soma, the middle part and distal part of the dendrite before HFS. The initial resting potential is −75 mV, however, with delivering the ongoing spontaneous activity to the granule cell, the new steady-state membrane potential of around −55 mV is maintained.

**Figure 3.3:**
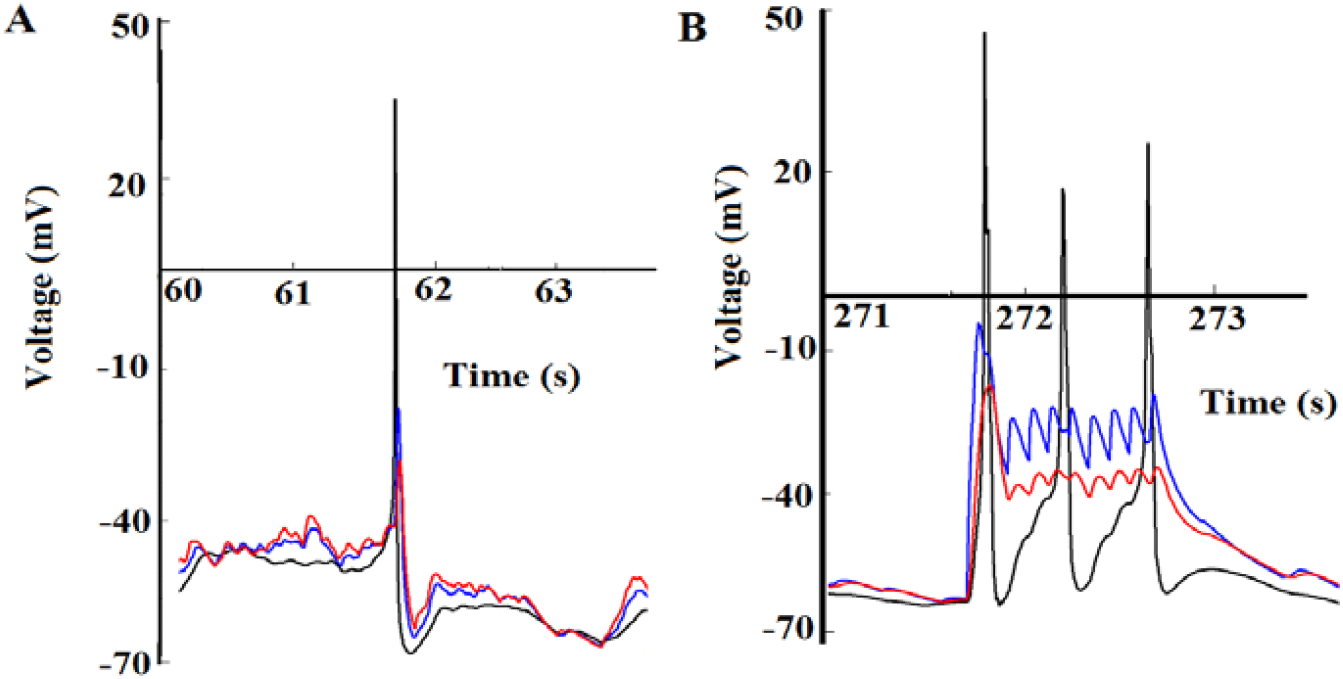
(A) Somatic membrane potential (black) and backpropagation of action potential to the distal (red) and middle (blue) parts of granule cell dendrites before HFS. New steady-state potential maintained by the ongoing presynaptic activity is −55 mV. The maximal after-hyperpolarization is −65 mV. (B) Somatic membrane potential (black) and backpropagation of action potential plus lateral and medial EPSPs to the distal (red) and middle (blue) parts of granule cell dendrites during one train of HFS.

When spatiotemporal summation of EPSPs evoked by spontaneous activity depolarizes the somatic membrane potential, the voltage increases to 30 mV. Backpropagation of action potentials to the middle and distal part of the granule cell depolarizes the membrane potential in the dendrites and raises the voltage in the synapses of the lateral and medial pathway to −23 mV and −33 mV respectively, which is above the dendritic threshold of −37 mV. Therefore, in both the lateral and medial synapses a postsynaptic event is detected, which in this case is a result of spontaneous input activity.

Figure 3.2B illustrates the increase of the EPSP of the granule cell during the first train of HFS delivered to the medial pathway. One train of HFS consists of ten input spikes at frequency 400 Hz. The summation of EPSPs evoked by the train of HFS depolarizes the somatic membrane potential and leads to the generation of a few action potentials (black traces). The backpropagation of action potential combined with EPSPs in the middle and distal parts of the dendrites is shown with the blue and red traces respectively.

As can be seen from Figure 3.3B, all medial postsynaptic events could pass the dendritic threshold of −37 mV. This shows that one train of presynaptic medial HFS can evoke ten post-events in the medial pathway (blue traces). Therefore, all 10 medial post events are paired with the preceding 10 pulses from the train of presynaptic (HFS) causing homosynaptic LTP to occur in the medial pathway. However, in the lateral pathway only five post-events (red traces) surpass the dendritic threshold while the others fail. In addition, presynaptic events are just spikes from the noisy spontaneous activity. Therefore, according to the ETDP rule, the five lateral post-events are paired with random spontaneous presynaptic spikes and result in synaptic depression rather than potentiation.

Another interesting phenomenon that we noticed in Figure 3.3B was the number of HFS-induced postsynaptic spikes and the difference in size of these spikes. Only 3 output spikes are evoked by 10 presynaptic pulses of each HFS train in the soma. Moreover, in this schematic picture, the first spike is the biggest one. We think the refractoriness property of the membrane mechanisms during each spike is the main reason for the subsequent spikes having a smaller amplitude than the first one. Refractoriness describes the property of a membrane, which dictates that immediately after the first action potential it is more difficult to generate a second spike (Gerstner, 2000). After the first spike, the membrane potential decreases below the previous potential of about −55 mV and becomes more negative. As we explained previously, at this point the membrane potential is in the after-hyperpolarization state of −65 mV, so that more stimulation is needed to increase the membrane potential and surpass the threshold for the next spike. Therefore, the second postsynaptic spike has a smaller amplitude than the first one. Moreover, the reduction in amplitude from the second to third spike is much smaller than the reduction in amplitude from the first to the second because of the increasing synaptic weight that occurs after the first spike.

#### 3.3.2 Floating average of postsynaptic activity

Figure 3.4A shows the membrane voltage when the first and the second burst of HFS are applied to the medial pathway. The first burst starts at time 275 s. One burst consists of 5 trains with the interburst interval of 1 second (each train consists of 10 pulses at 400 Hz) and the second burst starts at time 295 s. According to Figure 3.4A and B, when the membrane potential is mostly in the after-hyperpolarization status, the floating average of postsynaptic activity expressed by Equation (2.7) decreases. As can be seen in Figure 3.5, when the voltage of the membrane potential is between the initial values and after hyperpolarization values, the average of postsynaptic activity is in the lowest point.

**Figure 3.4:**
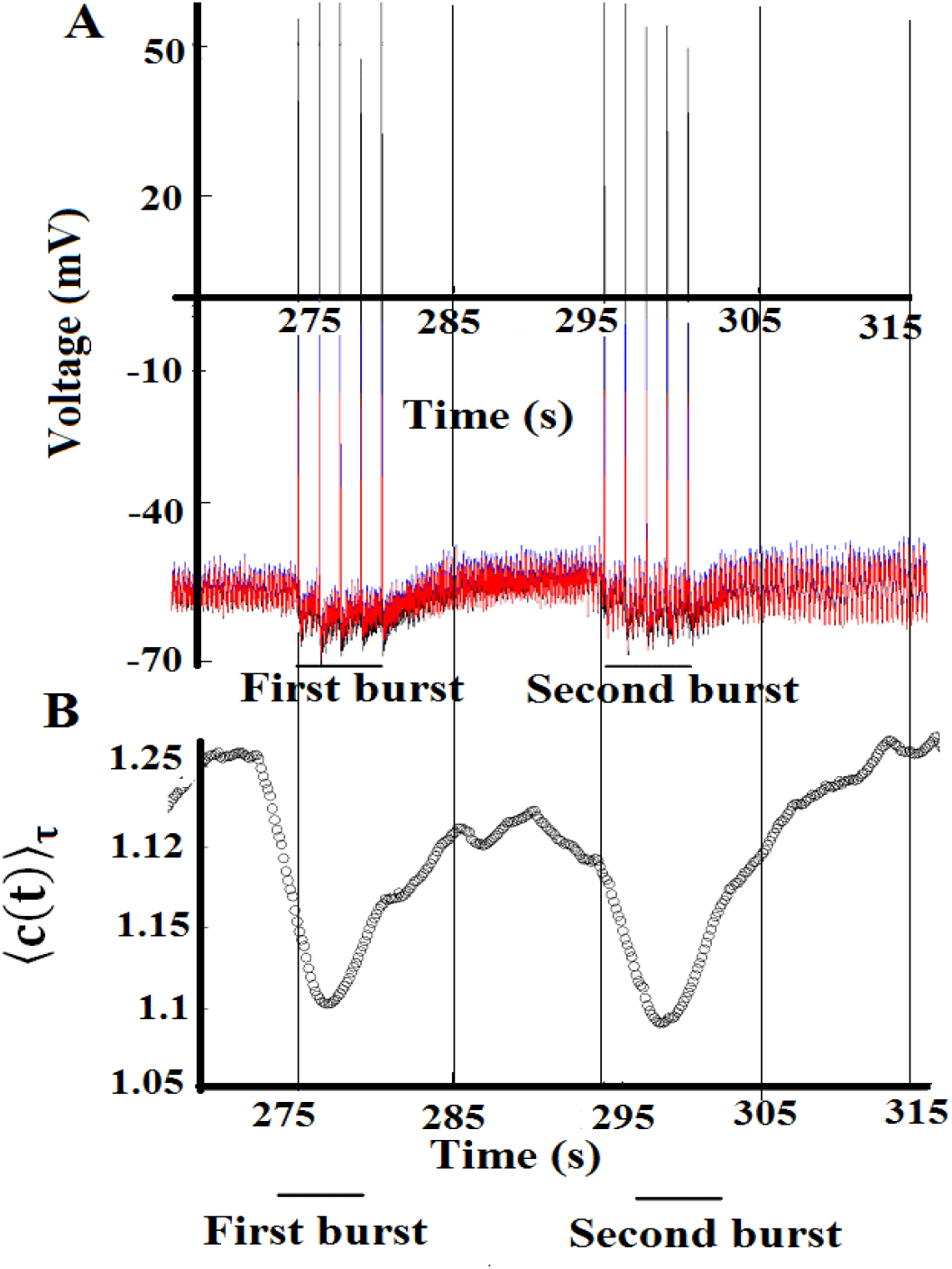
(A) Membrane potential just before HFS (when there is just spontaneous activity) and during the first and second bursts of HFS (in total there are 10 bursts). (B) Average of postsynaptic activity before HFS and during the first and second burst of HFS, with the exact timing as in A.

**Figure 3.5:**
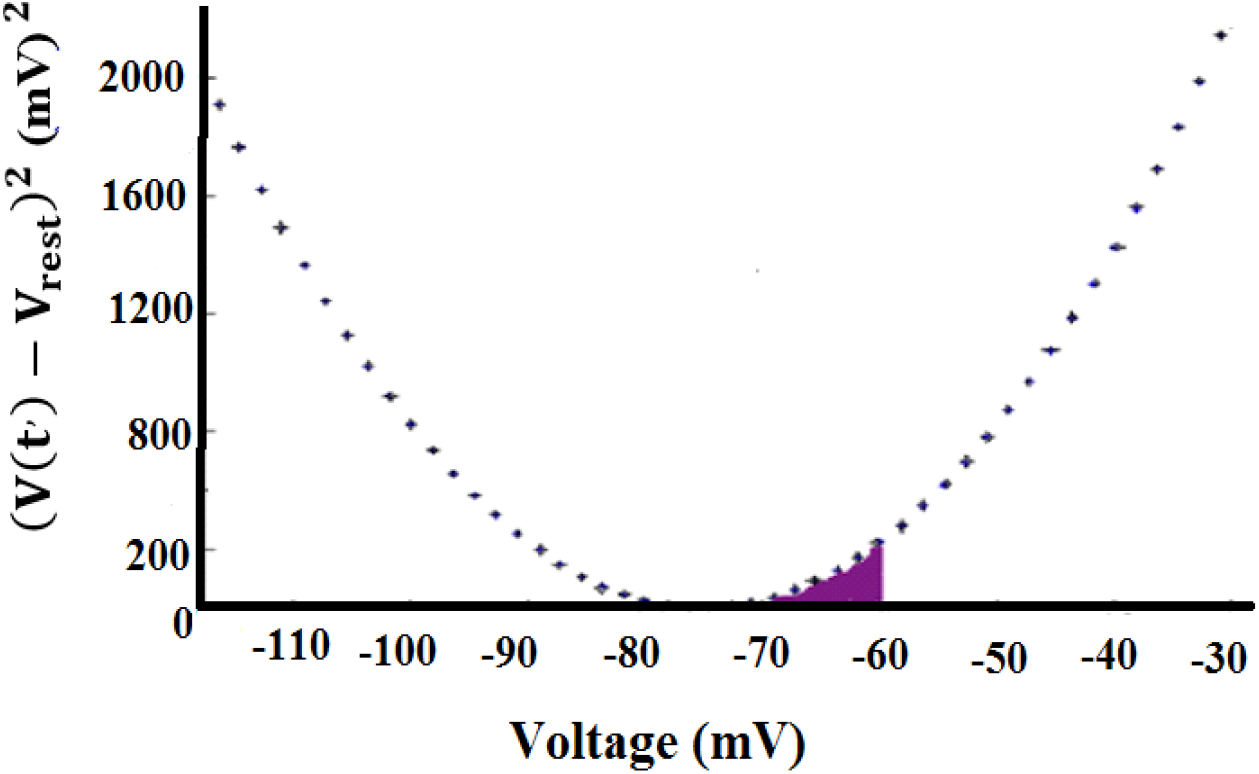
Parabola function (V (*t*’) – V_reset_)^2^ as a function of voltage.

Now we interpret the fluctuations in average postsynaptic activity. As the simulation starts, the average postsynaptic activity increases suddenly as the granule cell receives spontaneous activity. However, as the simulation continues, fluctuations in the average postsynaptic activity reduce until delivering the first burst of HFS. As can be seen from (Figure 3.4A and B) when this first burst of HFS is delivered to the medial pathway, the average of postsynaptic activity decreases because of the after-hyperpolarization feature of granule cell firing. During the time of 30 seconds between the two bursts of HFS, the average of postsynaptic activity again increases to the pre-HFS value because there is no tetanus. However, the reduction that occurs during the second burst of HFS is smaller than the reduction that occurs during the first HFS because the synaptic weights increased during this time as per the STDP rule. This scenario continues until all bursts are delivered. After HFS, because of the metaplasticity rule, the average of postsynaptic activity gradually reaches a steady state.

#### 3.3.3 Homosynaptic LTP and concurrent heterosynaptic LTD

To summarize, let us combine the nearest-neighbor ETDP and metaplasticity rules in explanation of homosynaptic LTP and concurrent heterosynaptic LTD. As the first burst of HFS is delivered to the medial pathway, the summation of EPSP and bAPs produce postsynaptic event that can pair causally with the medial HFS to increase the medial weight (ETDP rule). Also due to the decrease in the average postsynaptic voltage (Figure 3.4) the amplitude of potentiation P temporarily increases since P = P(0) / 〈c(t)〈_τ_, which also favours development of potentiation. However, during the 30-second gap between the first and second burst of HFS the medial synaptic weight does not change. When the next burst is delivered, the synaptic weight again increases and this sequence of events continues until the end of HFS. This synaptic potentiation phenomenon is called homosynaptic LTP (Figure 3.1). However, in the more distal lateral pathway fewer post-events surpass the dendritic threshold while the others fail. In addition, presynaptic events are just spikes from the noisy spontaneous activity. Therefore, according to the ETDP rule, the lateral post-events are paired with random spontaneous presynaptic spikes which results in synaptic depression rather than potentiation. Fewer postsynaptic events and constant D lead to LTD that has smaller magnitude than LTP. After HFS, because of the homeostatic nature of the metaplasticity rule, P returns to a pre-HFS value value and medial synapses stop keep growing, thus all the synaptic weights reach a new steady state.

#### 3.3.4 Necessity of metaplasticity

In this subsection, we examine whether the metaplasticity rules are necessary for inducing LTP and concurrent LTD in our granule cell model. For this purpose, we put P = P(0) and D = D(0) as constant values. As can be seen from Figure 3.6 we did not observe any LTP and concurrent LTD in this case. Therefore, we concluded that in our model metaplasticity should be involved in the ETDP rule in order to observe LTP and concurrent LTD.

**Figure 3.6:**
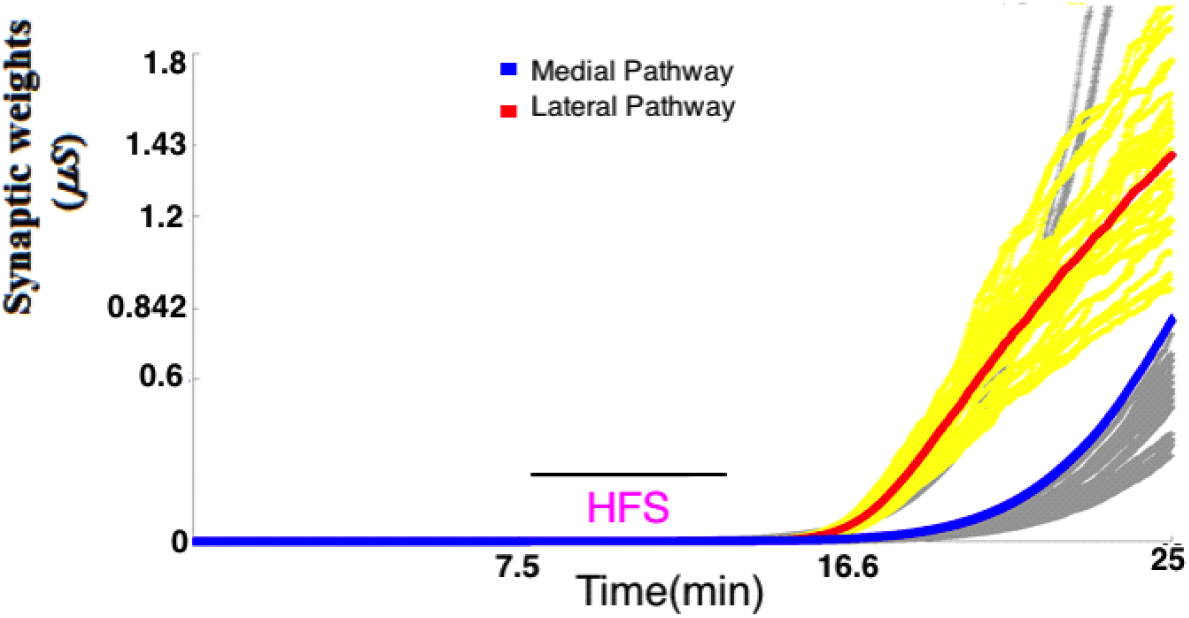
The average medial (blue curve) and lateral (red curve) synaptic weights with constant P and D.

### 3.2 Second simulation – no heterosynaptic plasticity when spontaneous lateral activity is blocked during a single HFS

In the second protocol to test whether or not the presence of noisy spontaneous activity is crucial for heterosynaptic LTD induction, we investigate the first part of the experimental studies from (Abraham et al.,2007) when a single HFS is applied to the medial pathway while lateral spontaneous activity is blocked during that time. In order to replicate the procaine inhibition of spontaneous lateral activity in experimental data, in our simulation, we switch off the lateral spontaneous activity during medial HFS. However, after HFS we switch the lateral activity back on again.

#### 3.4.1 Homosynaptic LTP and no concurrent heterosynaptic LTD

During medial HFS, the synaptic weights increase in the medial pathway (ETDP rule). After HFS, because of the ongoing medial spontaneous activity and the metaplasticity rule applied to the potentiation amplitude P, medial synaptic weights reach a steady state. Since we switch the lateral spontaneous activity off, there is no lateral activity in this pathway and thus no synaptic weight change as there are no presynaptic events and therefore no postsynaptic events. However, after HFS because we switch the lateral activity on again (albeit with the lower frequency because of the proposed long-term effect of procaine), the average lateral synaptic weight has almost no change or only a very small reduction (Figure 3.7).

**Figure 3.7:**
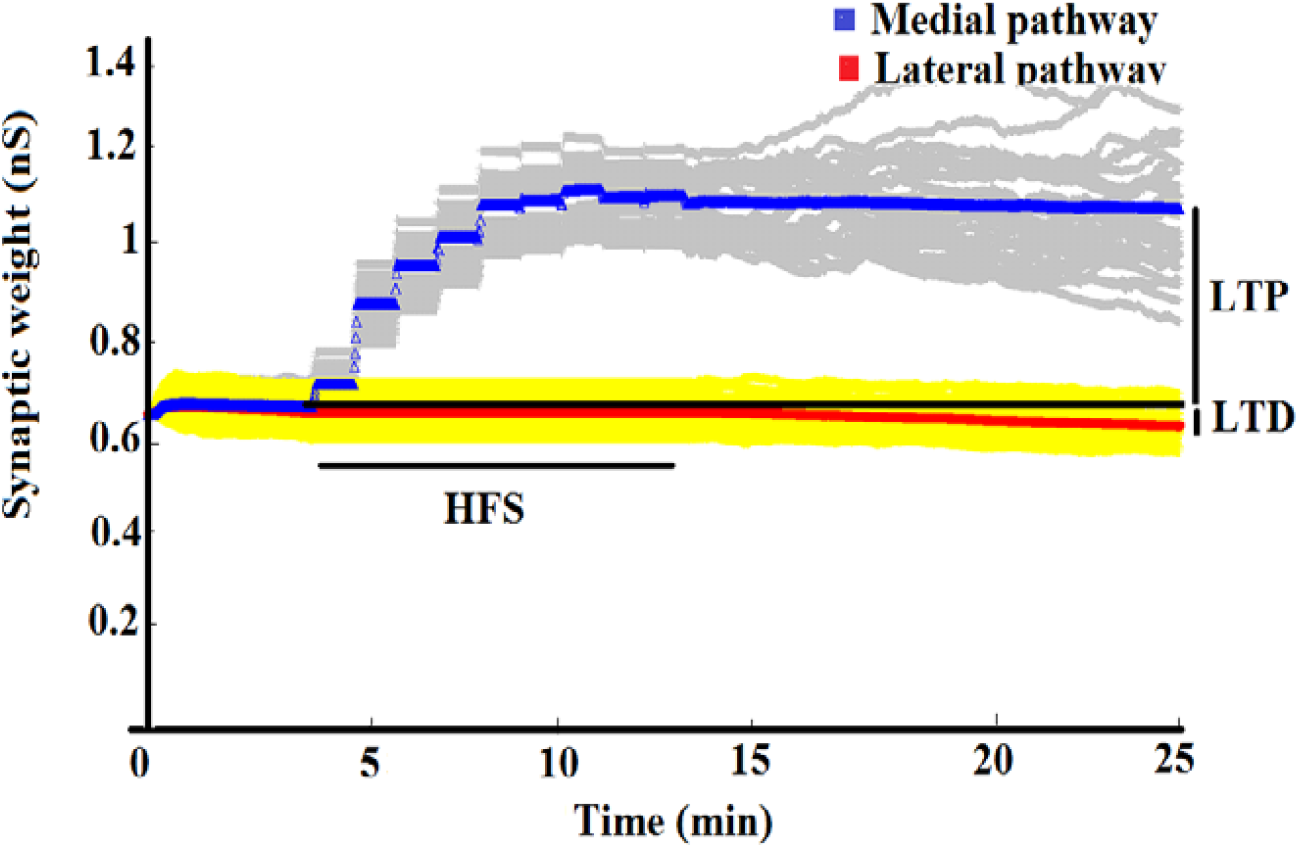
The blue curve is the average of medial synaptic weights and the grey curves are synaptic weights from 150 medial synapses. The red curve is the average of lateral synaptic weights and the yellow curves are lateral synaptic weights. As the HFS starts at the medial pathway, synaptic weights increase in this pathway. However, almost no LTD happens in the lateral pathway as we switch the lateral activity off during HFS.

In experimental studies, when procaine is injected to the lateral pathway, it takes about 60 minutes to wash out all procaine. We believe that during this period procaine may have a lasting effect on the level of spontaneous activity and decreases the frequency of the lateral activity. This means that after applying procaine it takes some time for the lateral activity to recover from the procaine inhibition and get back to the same frequency as before HFS. Therefore, we introduce a new parameter called lateral frequency after HFS (LAH) to characterize the frequency of the lateral activity after HFS, while other parameters are the same as given in Table 3.1.

Table 3.2 and Figure 3.8 show the magnitudes of medial LTP and lateral LTD as a function of lateral spontaneous frequency after HFS (LAH). The frequency of the lateral activity before HFS (during baseline) is 6.8 Hz. As the frequency of LAH is reduced, the size of LTD decreases as well. In the experimental data (Abraham et al., 2007), the LTP was 37±5% and LTD was −10 ± 6% after applying procaine. As can be seen from Table 3.2, when the frequency of the lateral activity after HFS (LAH) is 3.5 Hz we observe 60% LTP and −6% LTD compared to baseline. From this investigation, we concluded that to suppress LTD, the frequency of lateral activity after HFS should be less than 4 Hz to explain the procaine inhibition of heterosynaptic LTD with our model.

**Table 3.2.** Percentage of LTP and LTD as a function of frequency of lateral activity after HFS.

| LAH (Hz) | LTP (%) | LTD (%) | Postsynaptic firing rate (Hz) |
| --- | --- | --- | --- |
| 6.5 | 54 | -22.2 | 1.4 |
| 6.2 | 52 | -20 | 1.2 |
| 5.8 | 55.5 | -18.6 | 1.34 |
| 5.5 | 54.5 | -15 | 1.1 |
| 5.0 | 58.3 | -12 | 1 |
| 4.7 | 60.0 | -11 | 1.3 |
| 4.5 | 58.5 | -9 | 1.3 |
| 4.0 | 58.5 | -8 | 1.0 |
| 3.7 | 59.5 | -7 | 1.1 |
| 3.5 | 60.0 | -6 | 0.9 |

**Figure 3.8:**
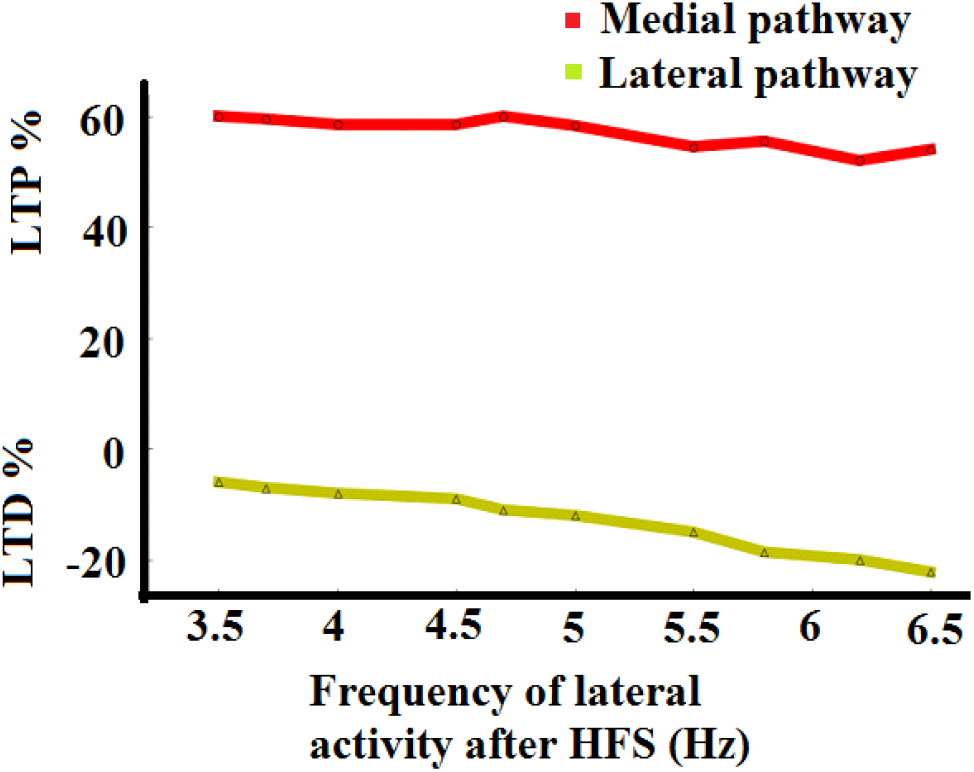
Amplitudes of LTP (red) and LTD (green) as a function of LAH.

#### 3.4.2 Average postsynaptic activity

The scaled running average of postsynaptic activity is shown in Figure 3.9. When the simulation starts and both pathways receive spontaneous activity, the average postsynaptic activity increases to 1.22 units. As the simulation continues, the average postsynaptic activity is stable. When we apply the first burst of HFS, it drops dramatically from 1.22 to 0.35. As we explained in the context of the first protocol, due to the after-hyperpolarisation after postsynaptic spikes during HFS the average voltage decreases. Moreover, in this case also the lateral activity is switched off during the medial HFS and the lateral pathway does not receive any input. Therefore, the average postsynaptic activity drops sharply. However, after HFS, because synaptic weights are increased in the medial pathway plus the lateral activity is switched on again, the average postsynaptic activity increases. As the simulation continues the average postsynaptic activity reaches a pre-HFS steady state.

**Figure 3.9:**
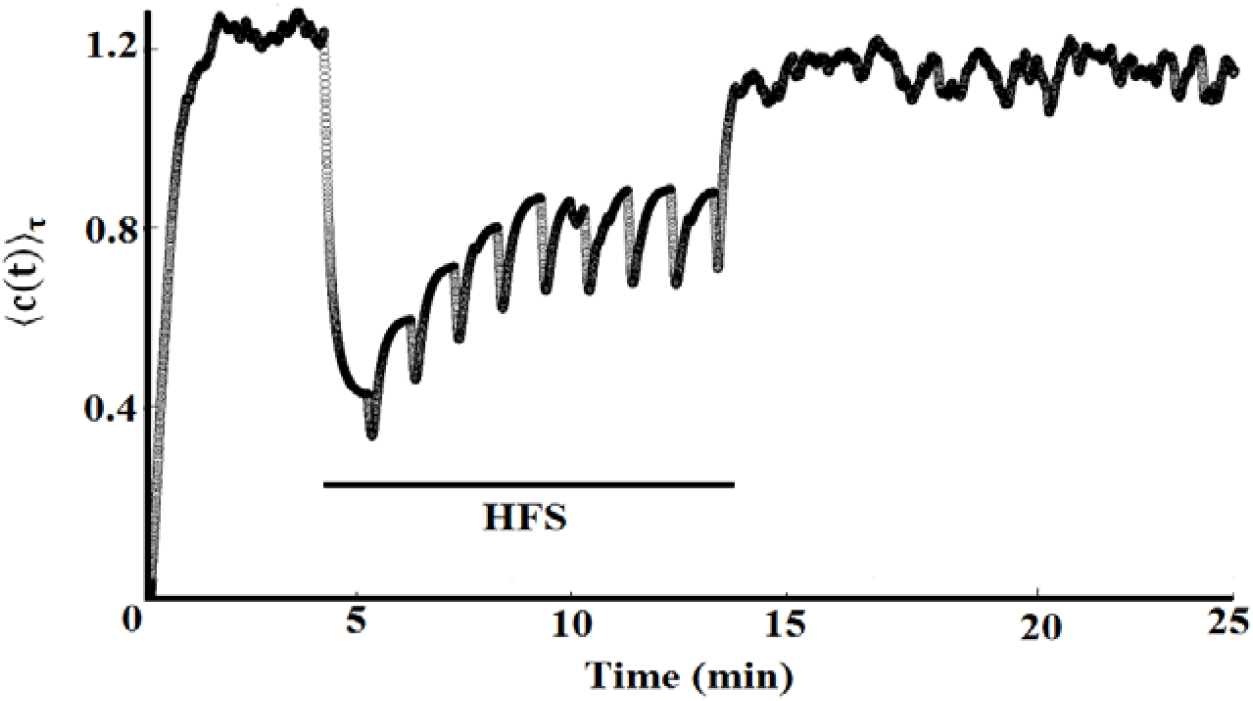
Scaled running average of postsynaptic activity before HFS is around 1.3. During HFS the average postsynaptic activity is very low with some oscillations because of HFS bursts and switching off the lateral activity. After HFS it rises again because of LTP and switching on the lateral activity.

### 3.3 Third simulation – effect of two consecutive HFS with all spontaneous activity being on

In this simulation as well as in the experiment (Abraham et al., 2007) two consecutive 400Hz DBS HFS were applied to the medial pathway. Simulation parameters are listed in Table 3.1. A few milliseconds after starting the simulation both medial and lateral synaptic weights and the average of the synaptic weights in the MPP and LPP become stable. As can be seen from Figure 3.10, when the first HFS is applied to the MPP the weight averages increase in this pathway and before applying the second HFS, 45% LTP is observed. As expected, in the LPP the average of the synaptic weights decreases and we observe −27% LTD. When the second HFS following the first one is applied to the MPP, the average synaptic weight slightly increases further in this pathway and we can observe a total of 54% LTP from the first baseline. This means we only get an additional 9% LTP from the second HFS. In the lateral pathway, we observe −39% LTD from the first HFS, and in this pathway, we also find only −12% more LTD because of the second HFS.

**Figure 3.10:**
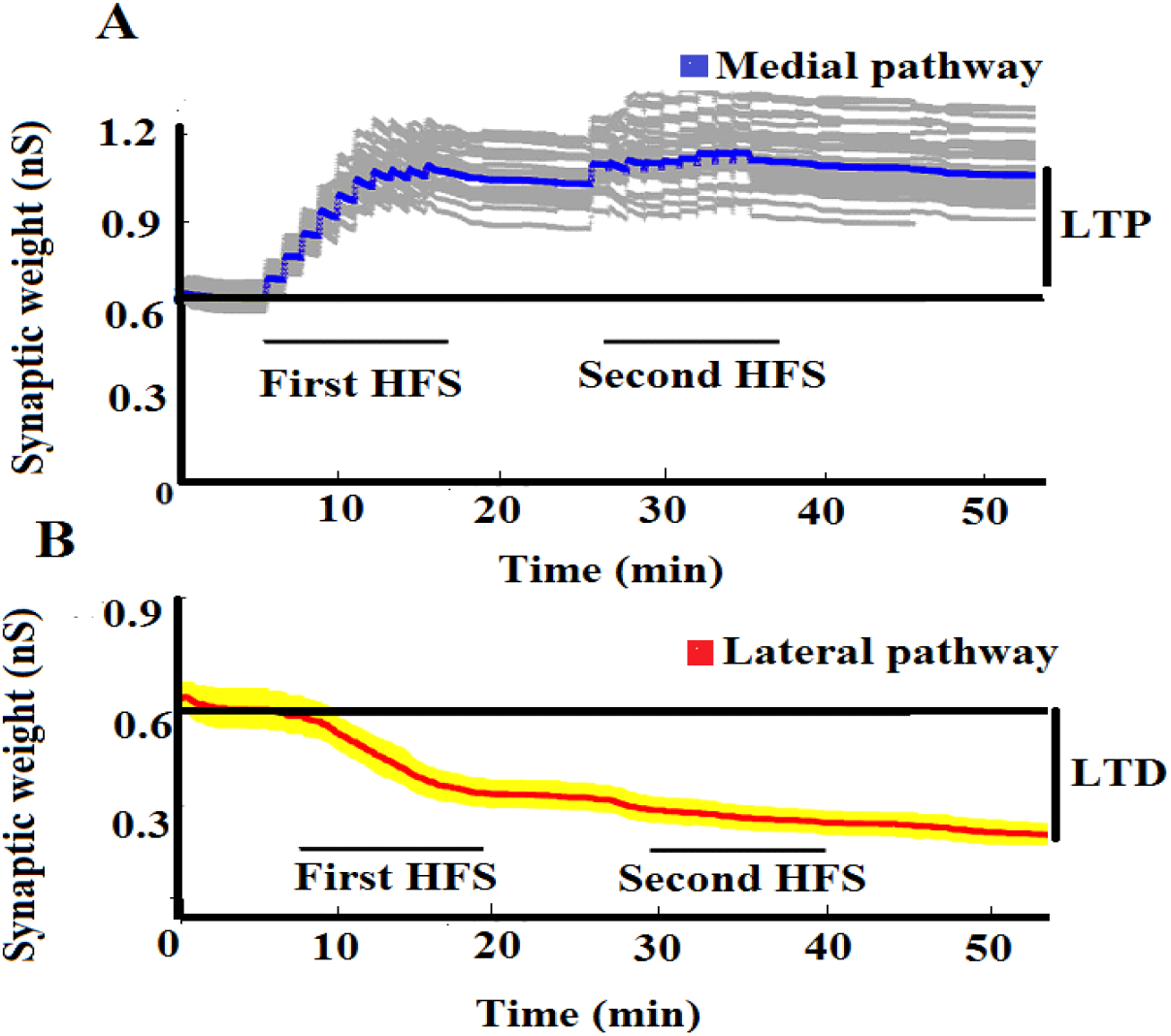
Results for two medial HFS with continued spontaneous activity being on in both MPP and LPP. (A) Individual medial weights (grey) and their average (blue). During the first HFS, the weight average increases up to 45%, and after applying the second HFS it increases slightly and 9% more of LTP is observed. (B) Individual lateral weights (yellow) and their average (red). After the first HFS, the weight average decreases by about −27%. After the second HFS, there is an additional decrease by −12%.

Metaplastic control of the potentiation amplitude P = P(0) / 〈c(t)〉–_τ_, due to the first LTP causes decrease in P, thus only a very small additional LTP is caused by the second HFS. Since the lateral spontaneous activity is on all the time, the heterosynaptic LTD occurs during both HFS.

### 3.4 Fourth simulation – effect of two consecutive HFS with lateral spontaneous activity being off during the first HFS

Before the first HFS, the average of both medial and lateral synaptic weights are stable. As can be seen from Figure 3.11, when the first HFS is applied to the MPP the weight average increases and 48% LTP is observed. However, in the lateral pathway because we switch the spontaneous activity off during the first medial HFS and then switch it on with the lower frequency than before HFS (3.5 Hz) as in the second simulation described in section 3.2, we only observe −6% LTD. With applying the second medial HFS, the average medial synaptic weight increases and we can observe a total of 56% LTP from the first baseline in the medial pathway. This means we get only an additional 8% LTP from the second HFS like in the previous protocol (section 3.3). However, in the LPP we observe only −10% LTD from the second HFS and in this pathway, we also find only an additional −4% LTD from the second HFS. Thus, reducing the post-procaine frequency of the lateral spontaneous activity has a lasting effect on blocking the induction of heterosynaptic LTD in the pathway.

**Figure 3.11:**
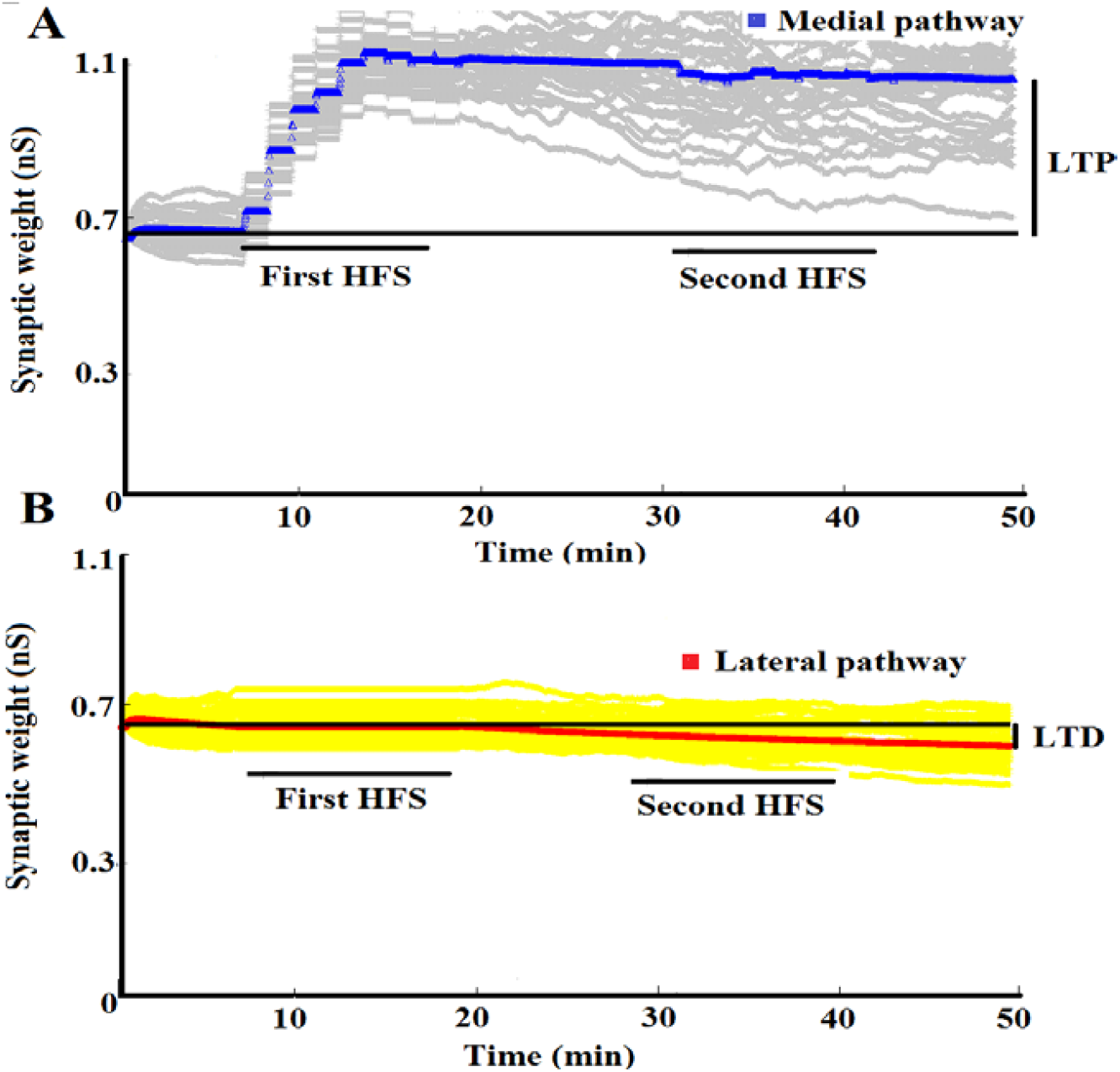
Results for two medial HFS and lateral spontaneous activity off during the first HFS (procaine simulation). (A) The blue curve is the average of medial synaptic weights and grey curves are the individual medial synaptic weights from 150 synapses. During the first HFS, the average of synaptic weights increases in the medial pathway but little further increase in synaptic weights occurs during the second HFS. (B) The red curve shows the average of lateral synaptic weights and yellow curves show the individual lateral weights from 150 synapses. In the lateral pathway as spontaneous lateral activity is off during the first HFS no LTD happens, while after the first HFS, as lateral activity switches on with the low frequency hardly any LTD occurs during and after the second HFS.

### 3.5 Reevaluation of the granule cell model

#### 3.5.1 The role of nine compartments on synaptic plasticity

Our granule cell model is based on nine compartments, four compartments for each dendrite and one for the soma. In this section, we assess the role of these compartments on the synaptic plasticity of our model. For simulations, we used the first protocol of the experimental studies from (Abraham et al., 2001). As the MPP transmits input to the middle dendrite of the GC, and the LPP transmits input to the distal dendrite of the GC (Scharfman, 2011), these compartments are necessary for our granule cell model. We also need the soma as the metaplasticity mechanism in our model needs the postsynaptic somatic voltage. We conducted a simulation without the proximal dendrite, and our observations showed that the firing rate before and after HFS was zero which means the neuron was silent. We think, as the proximal and middle dendrites are close, and the density of the fast sodium (Na) ion channels in the proximal dendrite is high, that our model, without proximal compartments, would not generate enough backpropagating spikes for LTP and LTD induction. We also ran the simulation without the granule cell layer (GCL). As the density of the Na channels in the GCL is high, we did not observe a match with experimental studies. Finally, we concluded that, at least for our plasticity model, all nine compartments are necessary.

#### 3.5.2 The role of ion channels on synaptic plasticity

The granule cell model introduced by Santhakumar et al. (2005) contains nine ion channels: fast sodium (Na), fast delayed rectifier potassium (fKDR), slow delayed rectifier potassium (sKDR), A-type potassium (KA), large conductance calcium and voltage-dependent potassium (BK), small conductance calcium-dependent potassium (SK) channels, and T-type (TCa), N-type (NCa), and L-type (LCa) calcium channels. These channel types are described by equations described in (Aradi and Holmes, 1999). The aim of this section is to evaluate the role of these ion channels on synaptic plasticity of the GC and investigate, which of these ion channels are necessary for synaptic plasticity in our model. For simulations, we used the first protocol of the experimental studies from (Abraham et al., 2001).

To assess the role of the fast Na channel on synaptic plasticity, we turned off this channel in all nine compartments. In our model, the amount of observed LTP was small and almost no LTD was observed at all. This finding supports the investigations from Jedlicka et al. (2015) that blocking the sodium channels in the dendrites interrupts heterosynaptic LTD. As the fast sodium channel regulates the amplitude and width of the action potential (AP) (Aradi and Holmes, 1999), when this channel is blocked, there is no action potential from the soma to backpropagate to the lateral pathway, which causes no LTD. Therefore, this ion channel is necessary for our plasticity model.

Next, to examine the role of the three potassium channels on synaptic plasticity, we turned them off in all compartments. Results showed that both LTP and LTD were not stable due to the action potential refractoriness which depends on the potassium channels (Jedlicka et al., 2015).

For the next simulation, we turned off the three calcium channels. Although we observed homosynaptic LTP in our model, the model was unable to produce stable heterosynaptic LTD and the results did not match the experimental studies similarly like in (Jedlicka et al., 2015). We think this is due to the role of these ion channels in regulating the spike back-propagation into the dendrites of the GC neuron.

Finally, we blocked the calcium-dependent potassium (SK) and (BK) channels. Without these ion channels, our model could replicate experimental studies. Therefore, our granule cell model does not include these channels.

## 4. Discussion

An important property of neural circuits is to maintain a homeostasis, by maintaining the same overall level of activity (Watt and Deasi, 2010). Experimentally observed properties show homosynaptic plasticity is not sufficient to stabilize the neural activity. Therefore, an additional mechanism is needed to keep the balance (Chistiakova et al., 2015). Thus, many experimental protocols that induce a homosynaptic change in stimulated synaptic weights also induce a heterosynaptic change in non-stimulated synaptic weights. In other words, during homosynaptic plasticity synapses need to be activated directly by presynaptic stimulation by experimental high-frequency protocol. However, for a heterosynaptic plasticity to occur, synapses do not need to be activated by experimental presynaptic stimulation for its induction (Chistiakova et al., 2014). According to several experimental studies, heterosynaptic plasticity mechanisms are necessary for complementing the homosynaptic plasticity during the regulation of neuronal activities (Chen et al., 2013).

In this work we simulate experimental studies, which show that if high-frequency stimulation (HFS) is applied to one input pathway of the dentate granule cells (GC) of freely moving rats or rats anesthetized with urethane, for instance the medial pathway, homosynaptic LTP appears in that pathway and simultaneous heterosynaptic LTD appears in the neighboring pathway, i.e., the lateral pathway and vice versa (Abraham et al., 2001; Doyère et al., 1997, Abraham et al., 2007). To model the Abraham’s et al. (2001; 2007) real experiments when HFS is applied to the medial pathway, we employed computational modeling with the reduced morphology multi-compartmental model of the GC. To model synaptic plasticity, we employed STDP-like rule, BCM-like metaplasticity rule, accompanied with the noisy spontaneous activity to model the heterosynaptic plasticity of the live dentate gyrus. The model setting is the same as in Jedlicka et al. (2015). Using the reduced morphology multi-compartmental model enabled us to introduce a local postsynaptic voltage threshold parameter for detecting the postsynaptic event, i.e. the supra-threshold local postsynaptic voltage, to be paired with the presynaptic spike arriving at the synapse. Thus, instead of STDP rule it would be more precise to speak about the ETDP rule, i.e. event-timing dependent plasticity in which the presynaptic spike and supra-threshold local postsynaptic voltage are events to be paired. The postsynaptic voltage crossing a given threshold can be the result of both – the backpropagating action potentials and spatio-temporal summation of EPSPs at the postsynaptic site of a synapse (Lisman and Spruston, 2005).

In addition, numerous experimental studies show that synaptic plasticity is a plastic phenomenon itself in the sense that the prior history of activity modifies the outcome of the current synaptic plasticity protocol (Abraham and Bear, 1996; Abraham 2008). This phenomenon is called metaplasticity or plasticity of synaptic plasticity, which is another mechanism that is involved to maintain neural homeostasis. Benuskova and Abraham (2007) and Jedlicka et al. (2015) proposed that the amplitude of potentiation P and amplitude of depression D in the ETDP rule are not constant but rather dynamically change their values as a function of the average of the postsynaptic activity over some recent past. In this paper, we introduce a modification, in which the running average of postsynaptic activity is calculated based on the difference between the postsynaptic voltage and resting potential at the soma (Equation 2.7). Besides the metaplasticity rule described in this paper, there are other metaplasticity rules such as those described by Clopath et al. (2010) and Zenke et al. (2013). In their metaplasticity rules, only the depression amplitude *D* depends on the average of postsynaptic activity while the potentiation amplitude is a constant value. Therefore, we were motivated by (Clopath et al., 2010) to find out, when only one of the potentiation or depression factors is subject to the metaplastic change, whether we can observe the same synaptic plasticity result. To answer this question, at first we ran our simulations with setting the *P* factor as a constant value and *D* as a function of the average of postsynaptic activity. We observed a good match compared with real experimental studies, provided we adjusted the scaling constant c0. We have also done our simulations with setting the *D* factor as a constant value and the *P* factor as a function of average of postsynaptic activity. After adjusting the scaling constant value c0, we can observe a good match with the real experimental data. We also ran our simulations when the *P* factor and *D* factor are both related to the average of postsynaptic activity. We surprisingly found, at least in our granule cell model, our synaptic plasticity results from either *P* or *D* factors being dependent on the average of postsynaptic activity are mostly the same as the results from when *P* and *D* factors both depend on the average of postsynaptic activity. Therefore, we concluded either potentiation and/or depression being dependent on 〈c〉_τ_ is enough to lead to homosynaptic LTP and heterosynaptic LTD phenomena in the model of granule cell. In this respect our results corroborate results of Jedlicka et al. (2015), but this time with the metaplasticity model based on the postsynaptic voltage instead of the postsynaptic spike count.

In our previous modeling work we demonstrated that one of the neural components of the hippocampus that controls the dynamics of homo- and hetero-synaptic plasticity is the noisy ongoing spontaneous activity in neural circuits (Benuskova and Abraham 2007; Abraham et al. 2007; Jedlicka et al. 2015). High-frequency stimulation (HFS) is applied to the medial pathway according to certain protocols (Abraham et al., 2001; 2007) and realistically simulated spontaneous activity is applied to both medial and lateral pathways during the whole experiments. In the tetanized medial pathway homosynaptic LTP occurs and simultaneously in the non-tetanized neighboring lateral pathway heterosynaptic LTD occurs. In addition to previous results, in the present paper, we investigated and showed that the level (frequency) of the lateral spontaneous activity determines the magnitude of the heterosynaptic LTD. The higher the lateral spontaneous activity, the larger is the magnitude of heterosynaptic LTD. Thus, no or low lateral spontaneous activity results in no or low heterosynaptic LTD. Thus for the first time, we replicated experimental results from Abraham et al. (2007), when lateral spontaneous activity was blocked and this blockade prevented induction of heterosynaptic LTD. In Abraham et al. (2007), HFS was again applied to the medial pathway. However, in this protocol, the experimenters blocked the spontaneous activity in the LPP by procaine during the duration of medial HFS. To simulate the inhibition effect of the procaine on the lateral pathway, simulated presynaptic spontaneous activity was switched off during the medial HFS. As in the real experiment, also in our simulations, no heterosynaptic LTD in the LPP accompanied medial LTP. We also modelled the effect of procaine inhibition of the lateral spontaneous activity during the first HFS upon the outcome of the second HFS. Thus, two HFS following each other were applied to the medial pathway except that the spontaneous activity was switched off in the LPP during the first medial HFS only. Then the lateral spontaneous activity was turned on again albeit with a reduced frequency. As before, no heterosynaptic LTD in the neighboring LPP occurred during the first HFS when LPP spontaneous activity was off altogether. Due to low frequency of the resumed lateral spontaneous activity, no heterosynaptic LTD occurred during the second HFS neither. The latter is however an assumption which remains to be experimentally tested.

To summarize, the goal of this paper was to model the homosynaptic LTP and concurrent heterosynaptic LTD in dentate gyrus granule cells (GC) *in vivo* using entirely voltage-based synaptic and metaplasticity rules. Using the reduced morphology multi-compartmental model of GC enabled us to introduce a local postsynaptic voltage threshold parameter for detecting the postsynaptic event to be paired with the presynaptic spike arriving at the synapse. The main novelty of this work compared to previous ones is that the metaplasticity rule is also made voltage-dependent instead of being dependent on the postsynaptic spike count over some recent past. Throughout this work we simulated an ongoing noisy spontaneous activity, which is a prominent characteristic of neural circuits *in vivo*. We varied the frequency of the lateral spontaneous activity in order to investigate that the level (frequency) of the lateral spontaneous activity determines the magnitude of the heterosynaptic LTD. We demonstrated that the higher the lateral spontaneous activity, the larger is the magnitude of heterosynaptic LTD. Thus, no or low lateral spontaneous activity results in no or low heterosynaptic LTD, which accounts for the results from Abraham et al. (2007), where experimenters for the first time showed that procaine inhibition of the lateral spontaneous activity during the medial HFS prevents development of the lateral heterosynaptic LTD *in vivo* for a prolonged time.

## Acknowledgments

This paper is based on the PhD Thesis of Azam Shirrafiardekani. We would like to thank W.C. Abraham for providing us with the original experimental data and many insightful discussions. Lubica Benuskova acknowledges financial support of the VEGA grant No. 1/0039/17.

